# Cell biological studies of ethanologenic bacterium *Zymomonas mobilis*

**DOI:** 10.1101/2020.07.20.213264

**Authors:** Katsuya Fuchino, Helena Chan, Ling Chin Hwang, Per Bruheim

## Abstract

Alphaproteobacterium *Zymomonas mobilis* exhibits extreme ethanologenic physiology, making this species a promising biofuel producer. Numerous studies have investigated its biology relevant to industrial applications and mostly at the population level. However, the organization of single cells in this industrially important, polyploid species has been largely uncharacterized.

In the present study, we characterized basic cellular behaviour of *Z. mobilis* strain Zm6 at a single cell level. We observed that growing *Z. mobilis* cells often divided at non mid-cell position, which contributed to variant cell size at birth. Yet, the cell size variance was regulated by a modulation of cell cycle span, mediated by a correlation of bacterial tubulin homologue FtsZ-ring accumulation with cell growth. The *Z. mobilis* culture also exhibited heterogeneous cellular DNA contents among individual cells, which might have been caused by asynchronous replication of chromosome that was not coordinated to cell growth. Furthermore, slightly angled divisions might have rendered temporary curvatures of attached *Z. mobilis* cells. Overall, the presented study uncovered a novel bacterial cell organization in *Z. mobilis*, the metabolism of which is not favoured for biosynthesis to build biomass.

**Importance:** With increasing environmental concerns about the exhausting use of fossil fuels, a development of sustainable biofuel production platform has been attracting significant public attention. Ethanologenic *Z. mobilis* species are endowed with an efficient ethanol-fermentation capacity that surpass, in several aspects, that of the baker’s yeast *Saccharomyces cerevisiae*, the most used microorganism for ethanol productions. For a development of *Z. mobilis* culture-based biorefinery, an investigation of its uncharacterized cell biology is important, because bacterial cellular organization and metabolism are closely associated with each other in a single cell compartment.

In addition, the current work highlights that polyploid bacterium *Z. mobilis* exhibits a distinctive mode of bacterial cell organization, reflecting its unique metabolism that do not prioritize incorporation of nutrients to cell growth. Thus, another significance of presented work is to advance our general understanding in the diversity of bacterial cell architecture.

## Introduction

Over the last two decades, there has been significant progress in uncovering the complexity of bacterial cellular organization (1). These developments significantly owe to recent advances in visualization techniques and genetic tools, enabling us to scrutinize bacterial cells. However, mostly due to the availability of tools, strains and established protocols, our knowledge of bacterial cellular organization is still limited to major model organisms such as *Escherichia coli*, *Bacillus subtilis*, *Caulobacter crescentus* and others. (1, 2)

It has been shown that several bacteria do not necessarily follow the canonical modes of growth and cell division displayed in model bacterial species (2–4). For example, species of the animal symbiont bacteria, *Thiosymbion*, divide longitudinally, exhibiting a striking contrast to the canonical vertical division (5). Polarized cell division in the intracellular pathogen *Chlamydia trachomatis* is another example of an unusual mode of propagation (6). Thus, although knowledge obtained from the bacterial models is immensely useful and covers common key mechanisms, it is unclear to what extent the studied mechanisms can be applied to other bacteria with specific metabolisms, physiologies or living strategies.

Facultative anaerobic bacterium *Zymomonas mobilis* is a natural ethanologen. Previous studies have extensively characterized its physiology with relation to industrial applications such as biofuel production (7). *Z. mobilis* is renowned for its relatively-simple and very fast catabolism, the Entner Doudoroff (ED) pathway coupled to the high activity of pyruvate carboxylase and alcohol dehydrogenase, which allows *Z. mobilis* to convert glucose to ethanol nearly at the theoretical yield under anaerobic condition (7, 8). Thus, only small amounts of substrate-carbon are incorporated as biomass in *Z. mobilis*, making this species an attractive biocatalyst in refinery systems.

Despite its industrially appealing physiology, its cellular organization has never been addressed in *Z. mobilis* biology. In contrast to other industrial workhorses such as *E coli*, *B. subtilis*, and *Saccharomyces cerevisiae* in which cell biology has been thoroughly scrutinized, only limited investigations have been done in *Z. mobilis* cells. This lack of understanding of *Z. mobilis* cell biology may be a bottleneck for manipulating its metabolism to be fully exploited, considering that cell growth and division are a consequence of the glycolysis that also produces ethanol as the major end-product. Thus, better understanding in the regulation of cell geometry is needed to guide rational metabolic engineering for *Z. mobilis* based biorefinery.

*Z. mobilis* belongs to Alphaproteobacterium, a bacterial group exhibiting a wide range of cell shapes and living strategies (9). Recent reports suggested that *Z. mobilis* cells deploy polyploidy (10, 11). Although polyploidy is prevalent across the kingdom of life, not much has been explored in bacteriology so far, as major model species are monoploids (12). Thus, investigating the *Z. mobilis* polyploidy and relevant cell biological features would deepen our understanding in the diverse repertoire of bacterial cell organization.

In the present study, we discovered and characterized the heterogeneous nature of *Z. mobilis* cells.

## Results

### Growth and division of single cell of *Z. mobilis* strain Zm6

Growth kinetics of *Z. mobilis* cells under various conditions are well-characterized at the population level (13). In contrast, single-cell growth of *Z. mobilis* has not been examined before, which might have overlooked important features of *Z. mobilis* biology.

We first determined cell size and shape of growing *Z. mobilis* strain Zm6 in the complex medium under anaerobic conditions and compared them with those of *E. coli* strain K12 aerobically growing in the LB medium. Microscopy imaging showed that cell volume of growing Zm6 cells were generally larger than those of growing *E. coli* K12 cells (Fig. 1A, 1B), the mean value of cellular length and width being 2.81 ± 0.71 μm and 1.28 ± 0.08 μm, respectively, for Zm6, and 2.16 ± 0.47 μm and 0.93 ± 0.06 μm, respectively, for K12. The Zm6 cells also exhibited a wider distribution of cell length compared to that of K12 cells (Fig. 1A, 1B). The Zm6 cells at later growth phase (OD_600_ = 1.5) in the same culture showed longer cell length (3.22 ± 0.75 μm, *n* = 315), possibly caused by the toxicity of accumulated ethanol in the culture. The doubling time for Zm6 cells in actively dividing phase was 105 minutes.

**Figure 1.**
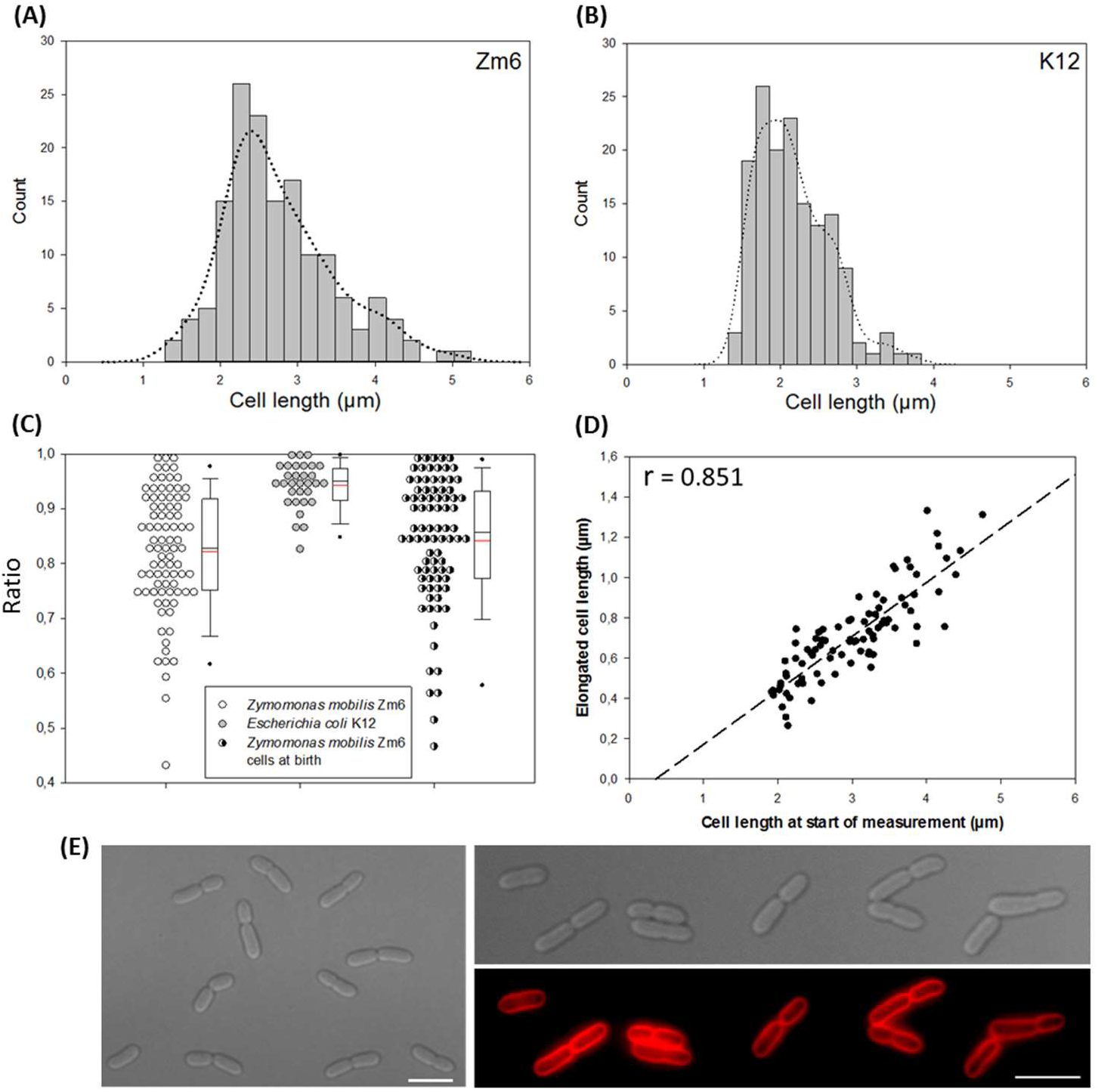
Cell size, morphology and single cell growth profile of *Z. mobilis* strain Zm6. (A and B) Histogram of cell length in growing *Z. mobilis* strain Zm6 (OD_600_ = 0.44, *n* = 150) and growing *E. coli* strain K12 (OD_600_ = 0.7, *n* = 150). Zm6 cell population exhibited a wide distribution in cell length than K12 cells. Kernel Density was plotted by a bandwidth being 0.25 for both histograms. (C) Dot density plot of cell length ratio between attached paired cell (length of short cell to long cell). Left plot; Zm6 pairs measured at random cell cycle stage (*n* = 85), middle plot; K12 pairs at random cell cycle stage (*n* = 31) right plot; Zm6 pair measured at its birth (*n* = 87). It was to be noted that there were not many attached-paired cells in the *E. coli* culture. Box-and-Whisker Plot were marked with mean (red) and median (blue). (D) A plot of elongated cell length during 40 minutes of growth by a function of cell length at the beginning of measurement. r; Pearson’s correlation coefficient, p < 0.001. *n* = 88. (E) Phase contract images of growing Zm6 strain (left and right top) and fluorescence image of Zm6 stained by membrane dye FM4-64 (right bottom). Zm6 was grown in the complex medium (OD_600_ = 1) under anaerobic condition and stained by Fm4-64 (20 μg/mL) for 15 minutes and washed by PBS before mounted on agarose pad for imaging. Note that attached two daughter cells often differed in its cell length from each other. Red; Fm4-64. Scale bar; 5 μm.

Zm6 cells formed rod-shape with elliptical ends (Fig. 1E). We also observed *Z. mobilis* two sibling cells that were often attached as a pair (Fig. 1E), as previously reported (14). Chained cells were also occasionally observed as well. Whilst, *E. coli* cells did not show such an attachment at septum in most of growing cells, implying that septal hydrolysing in *Z. mobilis* cells was not as active as in *E. coli* cells.

Interestingly, the attached-paired *Z. mobilis* cells often showed different cell lengths within a pair (Fig. 1E). We found this variant cell size between sibling cells intriguing, and further examined the cause and consequence of this variance.

An apical growth, the growth-mode employed by several alphaproteobacteria species (9), was a potential cause for the different lengths between sibling cells. To test if Zm6 cells grow apically, we stained the growing cells with Fluorescent D-amino acids to visualize sites of active peptidoglycan synthesis (15). The staining demonstrated that Zm6 cells grow at the septum and the lateral cell wall (Fig. S1), excluding an apical growth as the cause.

We then performed time-lapse imaging to monitor the growth of single cell for approximately two generations. Growing Zm6 cells were mounted on an agarose pad, sealed and left incubated for about 30 minutes before imaging. This was done to minimize microaerobic stress during the transfer, to ensure that Zm6 cells would continue to anaerobically grow during image acquisition. The series of imaging revealed that the constrictions took place often around, but not exactly at, mid-cell positions. Interestingly, division site selection was not strictly regulated to mid-cell position in Zm6 cells, exhibiting a sharp contrast to accurate mid-cell positioning found in many bacteria species.

Furthermore, when the cell length ratio of attached pairs at birth (short:long) was compared with the ratio of paired cells at random cell cycle stages, the mean ratio was higher in the pairs at birth (Fig. 1C). This indicates that another mechanism contributed to the different cell length within pairs during growth. We then measured elongation of cell length for 40 minutes in randomly selected Zm6 cells and plotted it as a function of cell length at the start of measurement. The plot showed a positive simple linear correlation (Fig. 1D), explaining that big cells grew faster than small cells, proportionally to cell size difference. In other words, the cell growth capacity per fixed cell volume was almost constant between individual cells.

The combination of two mechanisms, non mid-cell division and size dependent growth rate, could exert production of long, filamentous cells. However, we did not observe this. We therefore looked for what underlined here. By following the growth of paired sibling cells, we observed that longer cells divided earlier than the shorter ones (Fig. 2C). Among those paired cells with a length difference of more than 25% (i.e., short:long daughter cell length ratio being < 0.8), 74% of pairs (*n* = 27) showed that the longer cells started constriction 20 minutes earlier than its paired cell. To determine whether this was sibling-specific, we plotted cell cycle time vs. cell size at birth for individual cells, showing a negative regression (Fig. 2A). Hence, longer Zm6 cells tended to divide earlier than shorter ones, regardless of pairing. This was an intriguing paradox to us, having observed that the cells actively generated variant cell size over divisions, and at the same time, deployed a strategy to limit the size variance by modulating their cell cycle time.

**Figure 2.**
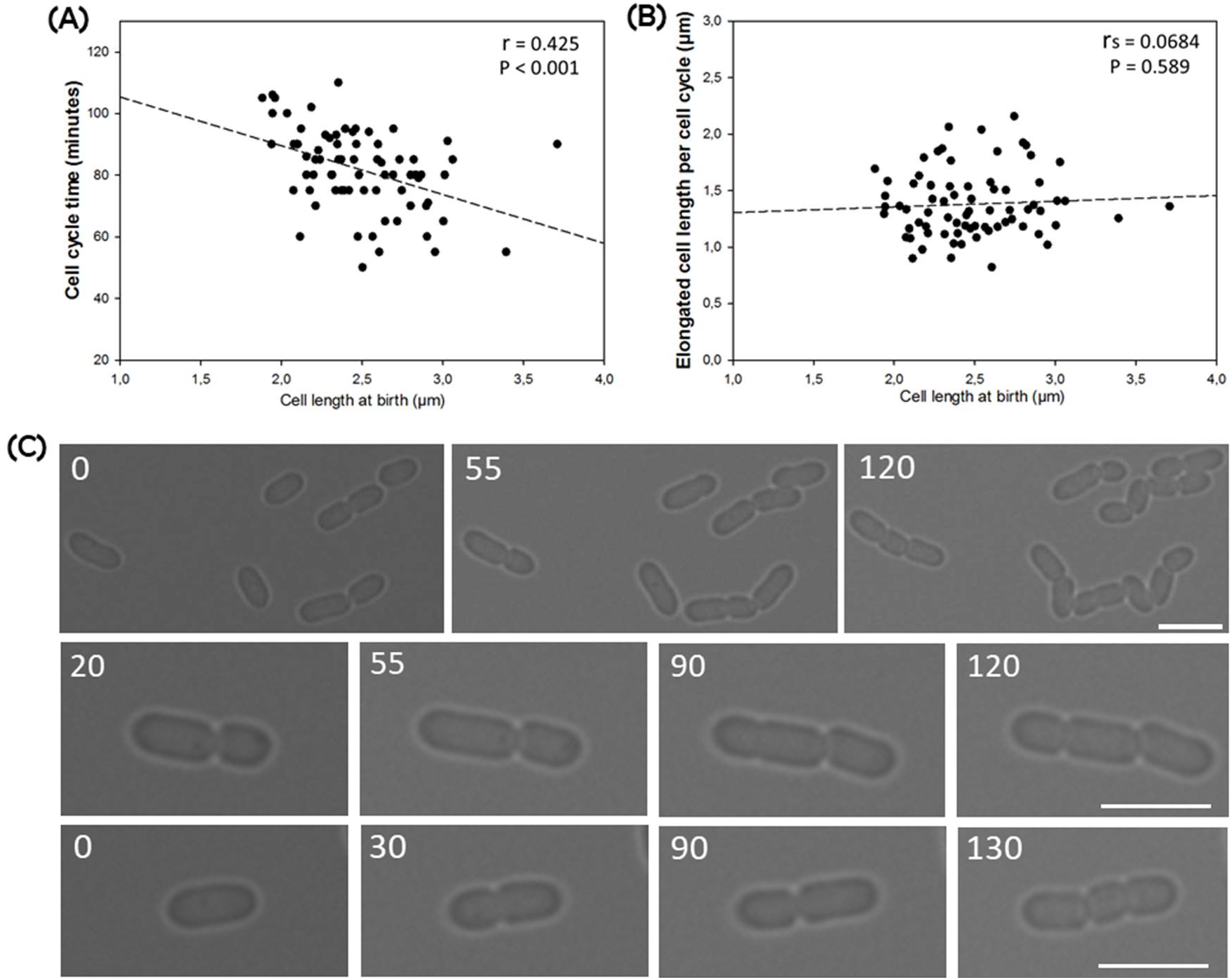
Time lapse imaging of growing Zm6 cells. Growing Zm6 cells in the liquid cultures (OD_600_ 0.5 – 0.75) were directly mounted on the complex medium agar pad and incubated for about 30 minutes before time-lapse imaging started. Cell growth and division were analysed using Image J (NIH). (A) Cell cycle time of individual cell was plotted against its cell size at birth. Negative correlation was shown by Pearson’s r, with p-value < 0.001, *n* = 73. (B) The plot of elongated cell length over one generation by a function of birth-size of measured cells. No correlation was found by Spearman’s rank correlation test, p = 0.589, *n* = 73. (C) Time-lapse imaging of growing Zm6 cells. Images are displayed over the time points, from left to right. Typical examples of an early division by long sibling cell is shown in the middle and bottom panels. Numbers indicate minutes after imaging started. Scale bars; 3.9 μm.

These observations reflected questions raised about bacterial size regulation, which has been a subject of intense debate (16–18). Several reports using single-cell quantitative approach have demonstrated that several bacterial species grow a fixed cell volume per cell cycle during steady-state growth, regardless of cell size at birth (19–23). The addition of a constant cell volume, described by the adder model (19–23), results in less cell size variation over cell cycles. However, the rule was not applied in *E. coli* under slow growth condition (22, 24) or in *Mycobacterium* species that grow asymmetrically by an apical growth (25).

We then attempted to examine if *Z. mobilis* cells applied the constant increment regulation. We measured ΔCL, the elongated cell length over a cell cycle in individual cells, and plotted it against its cell length at birth (Fig. 2A). It should be noted that several small cells generated during imaging did not complete division due to the time-limit of the method, and these were not included in the analysis.

The analysis displayed dispersed plotting, indicating that ΔCL appeared not to be strictly constant at individual cell level under our experimental setting. The relative standard deviation of ΔCL (21.2%) was significantly higher than those of other bacteria previously characterized (19, 26). The regression of plotting, however, did not show a slope over cell length, and no correlation was found (Fig. 2B). At population level, the cells may have been governed by the constant ΔCL regulations, despite that the governing principle was not tightly executed at individual cells. Further analysis using a more defined imaging method, combined with mathematical models, is required to identify the specific governing law of cell size homeostasis in *Z. mobilis*.

Taken together, the presented data provide an evidence that variant *Z. mobilis* cell size was generated by non mid-cell division, and further pronounced by size-dependent growth rates. The generated long cells executed division roughly at a timing according to added cell volume, or other unknown cues. Consequently, the short cells at birth have longer cell cycle span than long cells at birth. These cellular organizations appear to maintain variant cell size in *Z. mobilis* cell population, while preventing excessively sized cells.

### FtsZ sub-cellular localization in *Z. mobilis* cell

To gain further insights into *Z. mobilis* cell division, we fused *Z. mobilis* FtsZ (ZZ6_0455) to GFP to visualize its sub-cellular localization. Bacterial tubulin homologue FtsZ plays a central role in bacterial cytokinesis by recruiting proteins required for division (1). We constructed a pBBR plasmid derivative carrying *ftsZ*-*gfp* with the native promoter that allowed the expression in the introduced strain Z6KF2 (Fig. 3). Our analysis to examine the effect of the fusion (Fig. S2) led us to conclude that possible artefacts using the FtsZ-GFP is limited, and we used the merodiploid strain Z6KF2 for further investigations.

**Figure 3.**
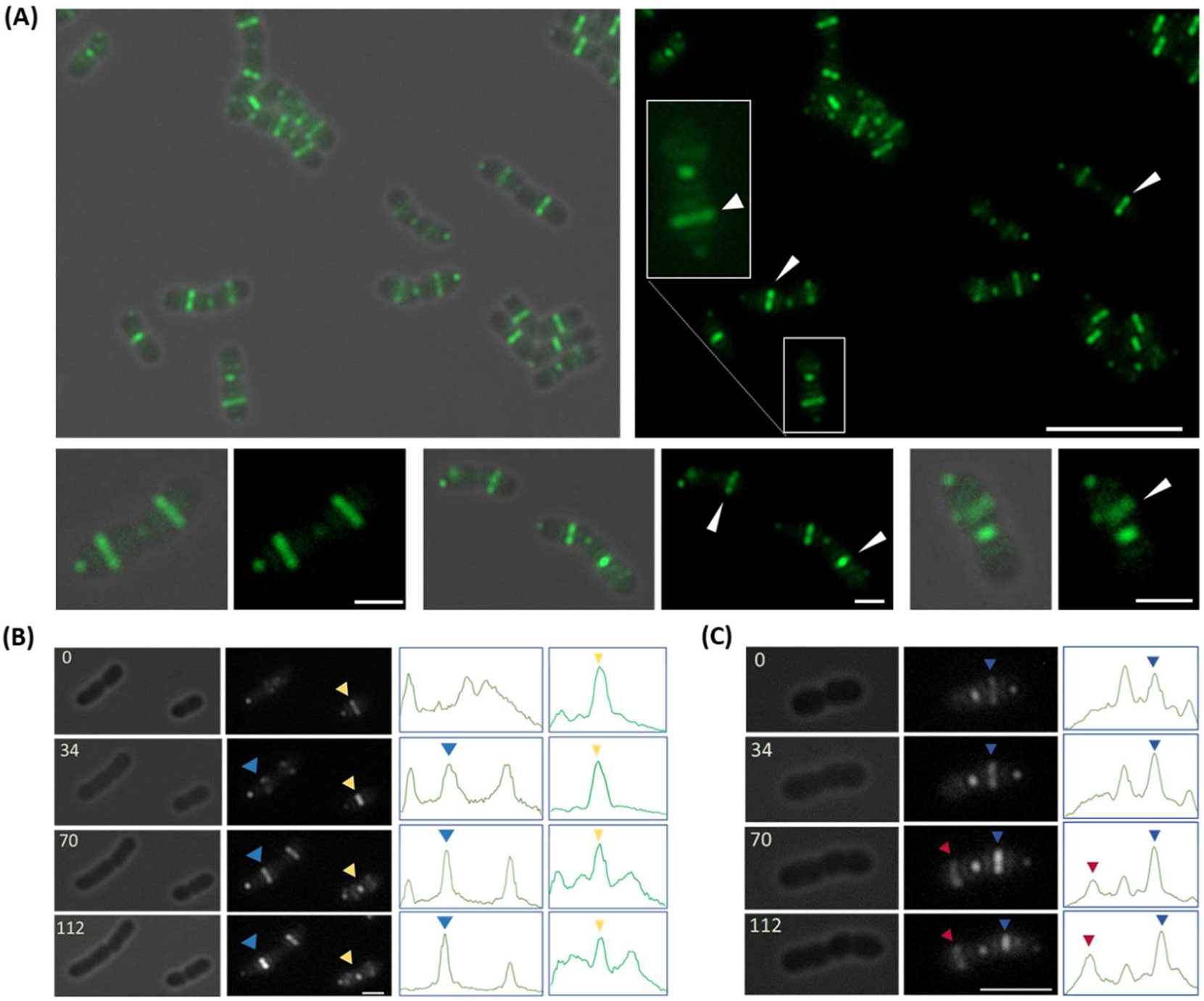
Sub-cellular localization of FtsZ-GFP in *Z. mobilis* strain Z6KF2. (A) FtsZ-GFP localizes around mid-cell as a Z-ring in many cells. Left, overlays of phase contrast and fluorescence images (green: FtsZ-GFP). Right, fluorescence images. Bottom panels show enlarged images of single cell. White arrowheads indicate cells that had earlier Z-ring formation or more abundant FtsZ in Z-ring in one of the paired cells. Scale bars; 10 μm for top panels, 2 μm for bottom panels. (B and C) Time-lapse imaging of growing Z6KF2 cells. Phase contrast image (left) was compared with fluorescence image (middle, white: FtsZ-GFP). The line profile (right) showed the relative FL intensities of GFP over long axis of each cell. Arrowheads colour indicates FtsZ-GFP fluorescence signal and corresponding peak in line profile. The numbers in phase contrast images indicate minutes after commencement of imaging. Earlier Z-ring formation was linked to earlier division in paired cells. Scale bars; 2 μm in (B) and 4 μm in (C).

Fluorescence imaging showed that FtsZ-GFP was often localized around mid-cell position as a ‘’Z-ring’’ in the merodiploid strain cell. FtsZ-GFP foci were also found at the free cell pole of many cells (Fig. 3), a phenomenon previously observed in other alphaproteobacteria (27–29).

Interestingly, many attached-pairs exhibited different FtsZ-GFP patterns within a pair; typically, a Z-ring had already formed in one cell while only crowds of signal were present in the other cell (Fig. 3A, 3C). In the case that both cells had already formed Z-ring, the fluorescence (FL) signal intensity of the Z-ring was generally higher in the larger sibling cell compared to the smaller cell (Fig. 3B).

A recent report suggests that FtsZ concentration is not constant per *E. coli* cell during cell cycle under slow growth conditions (30). The observed variant FtsZ abundance as a ring likely reflected different cell cycle stage of *Z. mobilis* cell. We found that 51.2 % (*n* = 82) of the paired cells showed more than1.5-fold difference of FL intensity peak of FtsZ-rings within a pair. Thus, temporal regulation of FtsZ expression often varied within paired cells, given that pairs were born at the same time. Therefore, not only its spatial regulation, but also temporal regulation of FtsZ was heterogeneous event among Zm6 cells.

Early Z-ring formation was possibly linked to early constriction in Zm6 cells, but polyploid bacterium might have checkpoints for chromosomal organizations that may delay the division. The Zm6 cells also divide at non-central positions, which might have a destabilizing effect to the Z-ring. We then performed time-lapse imaging to visualize FtsZ-GFP localization over the cell cycle. Imaging was performed at intervals of 15-20 minutes to minimize the effects of phototoxicity and bleaching. The imaging on attached-paired cells showed that formation of the Z-ring 20 minutes earlier in one of paired cells resulted in constriction 20 minutes earlier in that cell, for all observed pairs (*n* = 15, 100%). The data indicate that there were possibly no major check points to delay constriction after formation of Z-ring.

Furthermore, we found a positive correlation between FL peak intensity of FtsZ ring and cell size (Fig. S3). The plot shows that an FtsZ accumulation at the Z-ring generally increases with cell size, suggesting that large cells accumulate FtsZ at the Z-ring faster than smaller cells, taking into account faster growth rate of large cells (Fig. 1D). Thus, it appears that size or growth-correlated FtsZ-ring accumulation at division site is likely a key fundamental for temporal control of *Z. mobilis* division.

Interestingly, a recent study suggested that balanced biosynthesis and a threshold of accumulation of FtsZ are the key mechanisms for accomplishing the adder rule in *E. coli* cells at steady-state (21). Although it is not clear if *Z. mobilis* uses the constant increment rule for size regulation, the principle underlying *E. coli* adder rule may be applied in the size correction mechanism of *Z. mobilis* cells.

### Spatial FtsZ regulation in *Z. mobilis* cell

Next, to understand spatial regulation of *Z. mobilis* FtsZ, we made an *in silico* survey for *Z. mobilis* homologs of known cell cycle and FtsZ regulators in other bacteria. We found that minCDE complex and SlmA, prototypical negative regulators of FtsZ, are not encoded in the *Z. mobilis* genome (31, 32). Instead, the *Z. mobilis* Zm6 chromosome encodes *C. crescentus mipZ* homologue (ZZ6_0908). MipZ is a p-loop ATPase that negatively regulates FtsZ positioning in *C. cresentus* by being an antagonist of Z-ring formation (33). Notably, a recent report suggests that MipZ also regulates cell septation by localizing at mid-cell in *Rhodobacter sphaeroides*, alphaproteobacterim that carries two different chromosomes (34). Thus, *Z. mobilis* MipZ might possess an adapted function to the polyploidy, in addition to its function in the spatial regulation of FtsZ.

To gain additional insights into FtsZ spatial regulation, we set out to monitor the division process of stress-induced filamentous *Z mobilis* cell. A previous report demonstrated that Min proteins accurately regulate positioning of multiple Z-rings in the stress-induced filamentous *E. coli* cells, which assured cell size control of daughter cells (35). We exploited this type of stress response to visualize *Z. mobilis* FtsZ positioning regulations at a cellular level.

We first attempted a cephalexin treatment (40 μg/ml) to induce filamentations. However, we found that *Z. mobilis* has a high intrinsic tolerance to cephalexin, as *Z. mobilis* does to most commonly-used antibiotics (36). We sought an alternative approach and learned that the salt NaCl at a mild concentration induces filamentation and a bulge of single pole in *Z. mobilis* cells (37). Utilizing the response, we induced about 30 μm long filamentous Z6KF2 cell and monitored its FtsZ-GFP sub-cellular localization.

During the growth with saline stress (0.2 M NaCl), FtsZ-GFP formed foci or unclear structures in the filamented Z6KF2 cells (Fig. S4). After transferring the cells into the identical medium without salt, the FtsZ-GFP assemblies were reorganized to several ladder-like structures, as the cells started to regrow and divide (Fig. 4). During the recovery phase, we observed a randomized-fashion of FtsZ-GFP positioning, such as proximate double FtsZ-GFP rings (Fig. 4B, pointed by a yellow arrowhead at 80 minutes). Also, the timing of cell division was not coordinated with each other, as shown by simultaneous ladder-like divisions found in a part of cells (Fig. 4A, 152 minutes). The *Z. mobilis* FtsZ spatiotemporal regulation appeared not to be under tight control during the recovery, in contrast to one-by-one divisions organized by equally-spaced FtsZ-rings in the filamentous *E. coli* cells (35).

**Figure 4.**
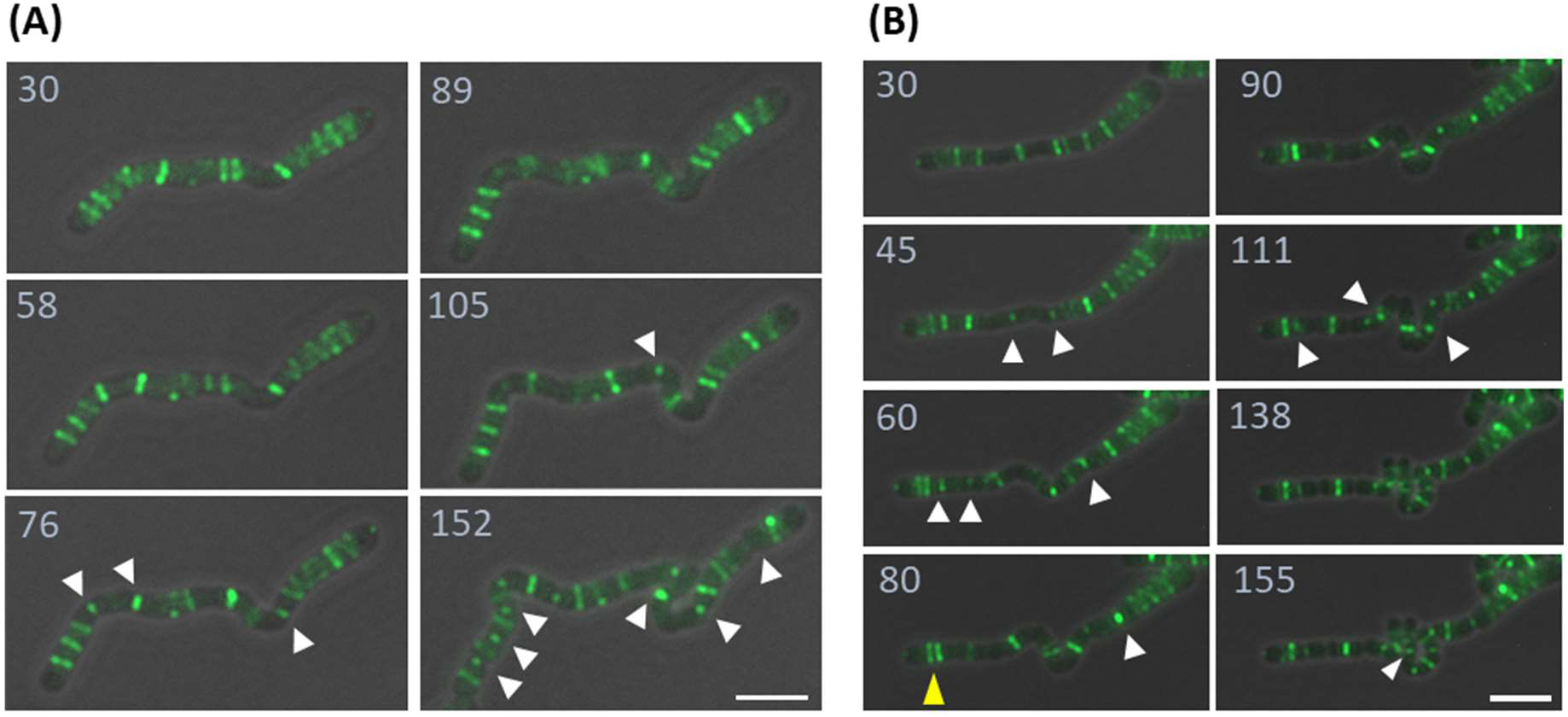
FtsZ-GFP sub-cellular localization in the filamentous Z6KF2 cells during the recovery from stress. Growing Zm6 cells were sub-cultured to the complex medium supplemented with NaCl 0.2M and grown for 5 hours. The cells were then transferred to an agarose pad of growth medium without additional salt. After 30 minutes of an incubation that allowed a recovery and re-growth of the transferred cells, imaging started. Time lapse imaging of two long filamentous cells were displayed as in (A) and (B). White arrowheads point to the positions where cell division was completing or just completed. Note that positioning of FtsZ-GFP was not tightly controlled in the filamentous Zm6 cells. In (A), a ladder-like FtsZ-GFP structure resulted in simultaneous divisions at 152 minutes. In (B), proximate double FtsZ-GFP-rings, pointed by yellow arrowhead, were observed. All images are overlays of phase contrast and fluorescence images (green: FtsZ-GFP). Numbers indicate minutes after cells were mounted on the complex medium agarose pad. Scale bars; 5 μm.

Importantly, the filamentous *Z. mobilis* cell managed to divide and regrow, in spite of the random positioning of division sites and timing, indicating that well-coordinated division was not necessary for cell growth. These observations implied that *Z. mobilis* might have evolved to produce different sizes of daughter cells during stress response, for better survival after recovery from stress.

### Chromosome organization in *Z. mobilis* cell

Previously, a transposon mutagenesis study using *Z. mobilis* strain Zm4 showed that the transposon-inserted mutants exhibited heterozygotes genotype, suggesting that *Z. mobilis* is a polyploid organism (11). The copy number of Zm4 chromosome (2.05 M bp) was estimated to be > 50, using qPCR (10). Based on similar genomic content between two *Z. mobilis* strains, we assume that Zm6 strain also possesses similar polyploidy feature. We therefore asked how multiple chromosomes organize in relation to growth and cell cycle in Zm6 cells.

We visualized nucleoids by staining live Zm6 cells with DNA-dye SYTO 9. The staining showed dispersed nucleoids localization throughout the cells, and no obvious condensation of nucleoid was found (Fig 5A). Interestingly, the averaged FL intensities within a cell compartment significantly varied between individual cells, implying that cellular DNA crowdedness or density was heterogeneous in the culture (Fig. 5A, 5B). We observed no anucleate Zm6 cells.

**Figure 5.**
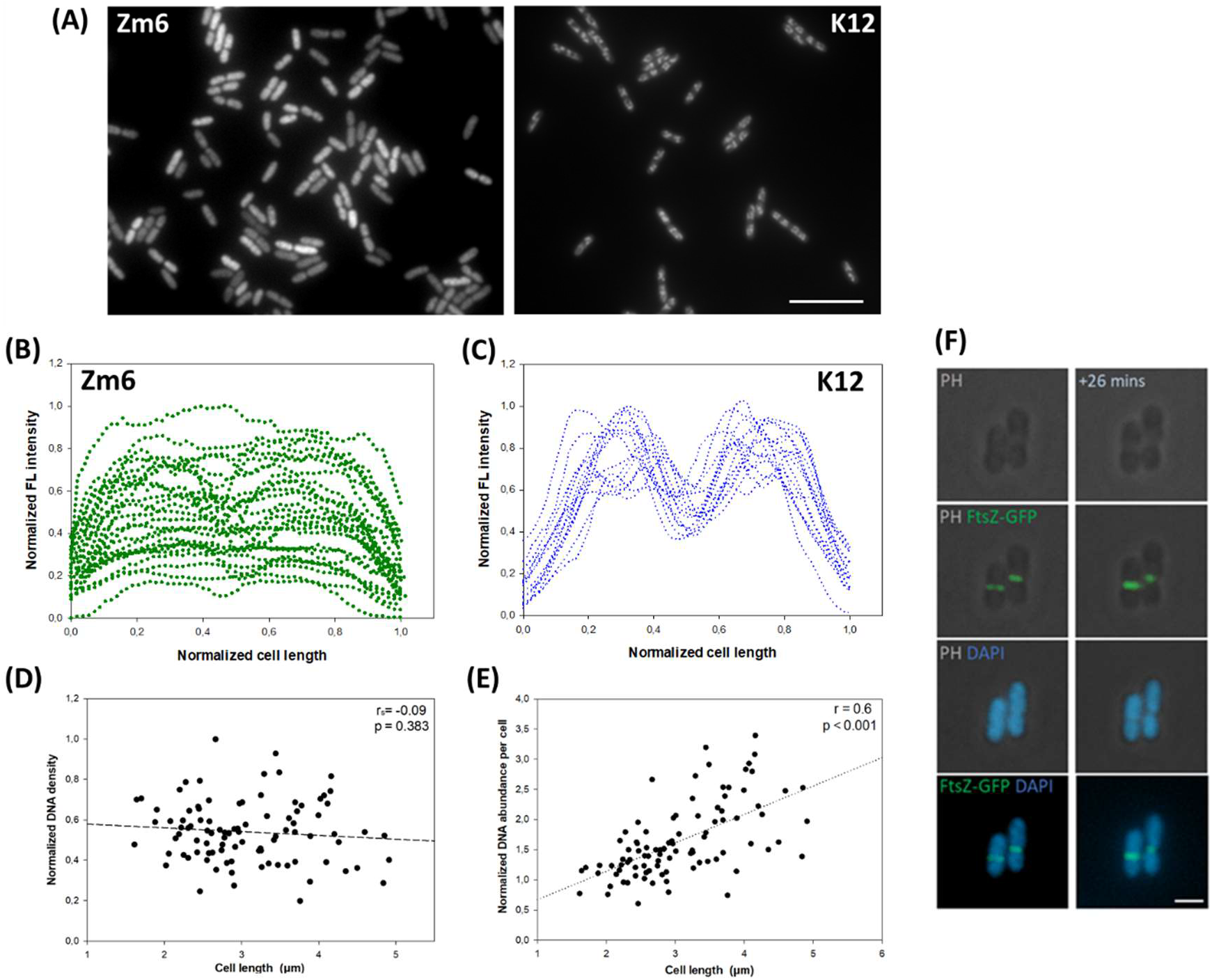
Analysis of stained nucleoids in *Z. mobilis* cells. (A) Fluorescence images of DNA staining in growing *Z. mobilis* strain Zm6 (left) and growing *E. coli* strain K12 (right). The Zm6 cells were stained by Syto 9 at 0.55 μM for 10 minutes. The K12 cells were stained by DAPI 5 μg/ml for 15 minutes. Both fluorescence signals are white in the images. Note that FL signal intensities varied between *Z. mobilis* cells. Scale bar; 10 μm. (B) FL signal profiling of 30 Zm6 cells over its long axis of each cell. FL signal and cell length were normalized. The profiles showed that the intensity varied between Zm6 cells. (C) FL signal profiling of 15 K12 cells over its long axis of each cell. FL signal and cell length were normalized. The K12 cells exhibited similar FL profiles between each other. (D) A plot of normalized average of FL intensity within the Zm6 cells, by a function of cell length of the measured cells. There was no correlation between two factors (Spearman’s rank correlation test; p = 0.589, *n* = 96). (E) A plot of normalized DNA abundance per cell compartment by the cell length of Zm6 cells. Positive correlation (Pearson’s r = 0.6) was found, *n* = 96. (F) Newly dividing Z6KF2 labelled with DAPI. Panels from top represent; phase contract images, overlay of phase contrast and fluorescence (green), overlay of phase contrast and fluorescence (blue), overlay of two fluorescence (green and blue). Right images were taken 26 minutes after left images. Green; FtsZ-GFP, Blue; DAPI. Scale bar; 2 μm.

For a comparison, we stained *E. coli* strain K12 by DNA-dye DAPI and observed the similar localization patterns of *E. coli* nucleoid to those previously reported (38, 39). Nucleoids occlusions and condensations were observed in all the *E. coli* cells (Fig. 5A). The intensity of signal was found at relatively the same level between different K12 cells (Fig. 5A, 5C).

We initially suspected that the variant signal intensities in Zm6 cells might be an artefact from the staining procedure. We therefore attempted to exclude the possibility by employing different staining methods, and similar results were obtained (Fig. S5). As *Z. mobilis* is a polyploid, we concluded that the growing Zm6 culture exhibited heterogeneous cellular DNA crowdedness/density among individual cells.

Interestingly, up to 1-3 fold FL intensity variance was found between similar-sized Zm6 cells (Fig. 5D). Non mid-cell division was a potential suspect for causing uneven distribution of macromolecules to daughter *Z. mobilis* cells. Nevertheless, most of the attached sibling cells, including newly dividing pairs, showed very similar FL intensity level within a pair (Fig. 5F, S6). This demonstrated that division was not the main cause for the large intensity variance.

Here, another interpretation of the equal signal intensities within pair is that chromosome numbers per single cell compartment varied between sibling cells, roughly proportional to cell size difference (Fig. 1C). Thus, interestingly, the multiple copies of chromosomes in parental cells appeared to be unequally distributed to the daughter cells, according to their birth-sizes, to ensure that chromosome numbers per cell volume, DNA crowdedness, became roughly equal between daughter cells. This type of chromosome distribution resembles the observation made in the *minD* mutant of polyploid cyanobacterium *Synechococcus elongatus* (40), and there may be a conserved mechanism in multiple chromosomes spatial organization among the polyploid bacteria. Yet, the *S. elongatus minD* strain produced anucleate cells, while *Z. mobilis* zm6 strain did not produce such cells.

If not by non mid-cell divisions, what would cause the variant DNA crowdedness between Zm6 cells? DNA crowdedness within a cell was found not to correlate with its cell length in the Zm6 culture (Fig. 5D). Yet, DNA abundance per cell compartment (averaged FL intensity integrated over the cell length) showed a modest positive correlation with cell size (Fig. 5E). This correlation resembles cell size to-genome copy number coordination in the polyploid bacterial species. For example, step-wise, asynchronous replication was found in *S. elongatus* to ensure a positive correlation between chromosome numbers and cell volume (40, 41). Similar correlation was found in the symbiont bacterium *Sinorhizobium meliloti* during its differentiation to bacteroid (42, 43) and the gigantic bacterium *Epulopiscium* species (44). Thus, asynchronous chromosome replication system that coordinated with cell volume expansion may be a shared trait among bacteria with multiple copies of same chromosome.

In this context, we speculate that asynchronous chromosome replication might have been loosely coordinated with cell growth in *Z. mobilis*, and the miscoordination may have caused the heterogeneous DNA crowdedness over generations. However, this should be experimentally verified by live imaging of *Z. mobilis* chromosome numbers in growing cells.

Overall, our presented data suggestively argue that polyploid bacterium *Z. mobilis* possesses a novel mode of chromosome organization. The heterogeneity of cellular DNA crowdedness observed in *Z. mobilis* culture appear to be a unique arrangement even among polyploid bacteria, because the correlations of cell size with DNA contents were found to fit well in the polyploid bacteria species such as *S. elongatus* and *Epulopiscium* species (44, 45), indicating that these species do not exhibit cellular heterogeneity of DNA crowdedness observed in *Z. mobilis*.

### Cell curvature

Growing Zm6 cells often existed as attached pairs (Fig. 1), constituting 36 % of growing population (OD_600_ 0.8, *n* = 430). It was eye-catching that, in some attached-pairs, an extended long cell-axis from the daughter cells were crossing with each other, instead of in line with each other (Fig. 1E and Fig. 6A). In other words, a rim shape of two attached-cells is curved or bent, despite the semi-symmetric shape of each single cell.

**Figure 6.**
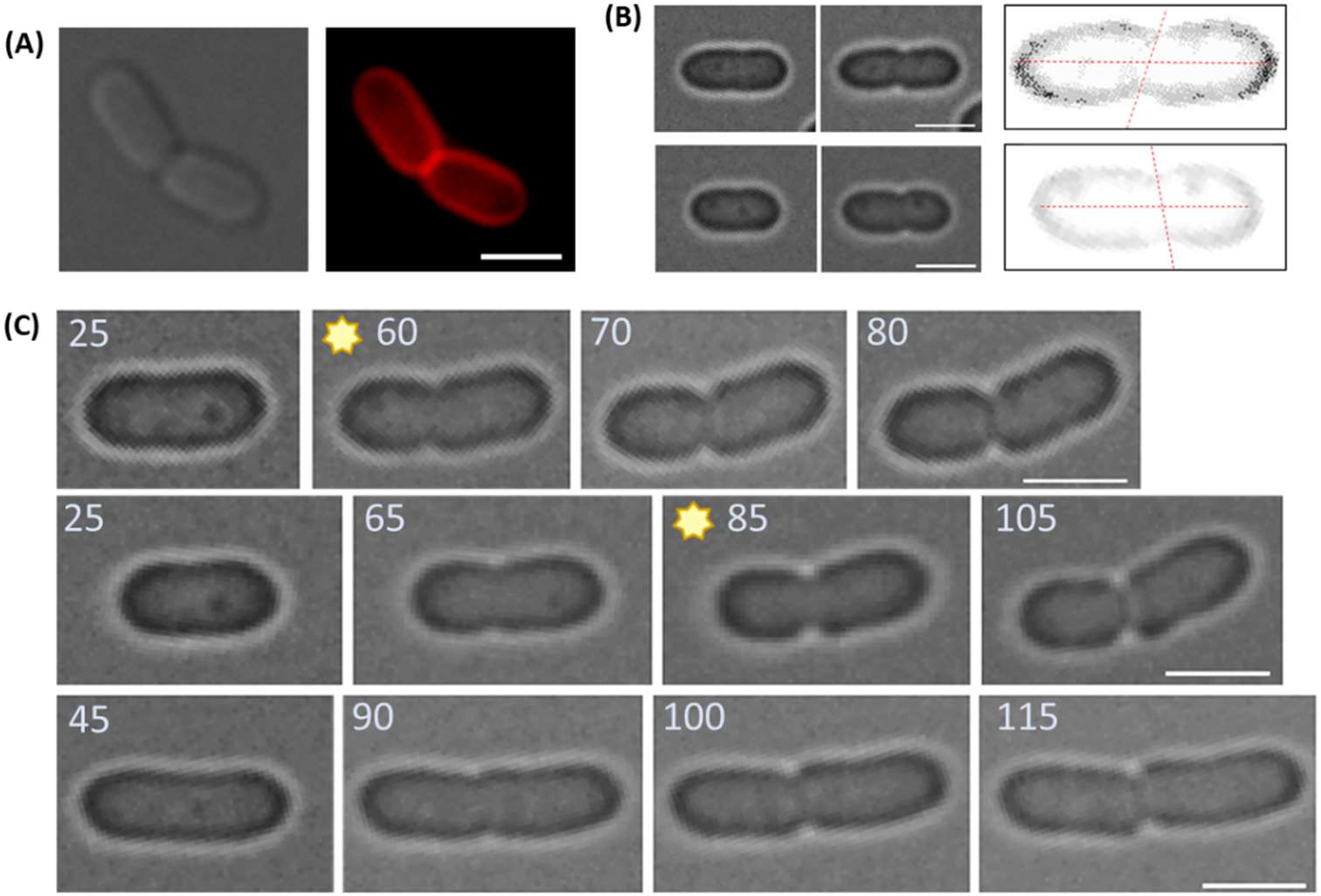
Curvature of attached *Z. mobilis* sibling cells. (A) The fm4-64 labelling on sibling Zm6 cells that were curvedly attached. The images are phase contrast image (Left) and fluorescence image (right). Red; Fm4-64. (B) Angled division in Zm6 cells. Phase contrast images of Zm6 cells before (left) and after (middle) constrictions took place. In the right image, a rim of cell was pronounced by adjusting the contrast of image. Two red dotted lines, showing the constriction axis and the long axis of the cell, show that the division is not 90 degree angled to the long axis of cell. (C) Time-lapse imaging of single Zm6 cells showing the oblique-like division. Division of three cells are separately displayed in time order (from left to right). The numbers indicate minutes after a commencement of imaging. The images highlighted with star sign shows the curved cell before a completion of division. Note that cells in upper two panels showed the attached-curvature, while the cell in the bottom panel did not generate the curvature despite the inclined division. Scale bars; 2 μm in all images.

These joint-curved cells were fractional in growing culture, constituting 10.4 % of the total population (pairs with an angle between long axis > 20°, *n* = 430). The Zm6 cells were not separated right after divisions, likely due to temporary insufficient activity of peptidoglycan hydrolase, but how attachment-based curvature was generated?

Uneven cell-wall hydrolysis activity could be one cause of the curvature, but it might require an extreme delicate control. Another possible cause was that the curvature was mechanically introduced during the division process. With this in mind, we observed that constrictions in dividing Zm6 cells were often found slightly inclined at the short axis of cells, i.e., the constriction occurred at a slightly oblique angle (Fig. 6B). For clarification of the angled division, we imaged electron micrographs of *Z. mobilis* cells. The images suggested that constriction was slightly inclined in some of *Z. mobilis* cells (Fig. S7).

We then followed the fate of cells that divided in a slight oblique manner. We noticed that the curvature/bent of a single cell was occasionally generated upon the slanted constriction, but before completing the septum closure. (Fig. 6C, panels highlighted with star). When the division was completed, the two attached cells exhibited the mentioned curvature (Fig. 6A and 6C). When the cells divided without an obvious angle from the short axis of cell, the bent of cell was not observed (Fig. 1C, bottom panels). These observations led us to speculate that an angled division might be a cause of the observed curvature of attached cells.

However, it should be noted that the timing of separation of attached cells appeared to be random among *Z. mobilis* cells. A collision of two separated elongating daughter cells, regardless of angle of division that just happened, could result in a similar appearance to the attached-curve during time-lapse imaging. This was the source of our initial scepticism to the speculation. Yet, the collision of cells eventually resulted in sliding of cells, which was distinguishable from the attached-curved cells. The curvature was also observed in chemically-fixed cells as well, in which colliding cells were excluded in imaging. More importantly, the curvature of single cell was generated before the completion of division (Fig. 6C), suggesting that the dividing process by itself could generate the curve by altering the vector of turgor in cells.

Also, it should be pointed out that an angled constriction did not always introduce attached-curvatures to cells. The case without generation of curvature is as shown in the bottom panels of Fig. 6C. Our observations suggest that combination of non-mid cell division and angled constriction led to the curvature, but not either of the mechanism alone did. Further examinations on mechanical property of *Z. mobilis* cells in relation to positioning and angle of divisions is required to understand the mechanism of curve generation.

## Discussion

*Z. mobilis* is well-known for its extremely efficient homo-ethanol fermentation, which may have overshadowed other unique aspects of *Z. mobilis* biology. For example, its aerobic respiration yields very little or no energy (8, 46). *Z. mobilis* carries a complete electron transport chain (ETS), which is constitutively expressed under an anaerobic condition (8, 46, 47). Despite its high rate of oxygen respiration surpassing that by *E. coli*, and possession of F_0_F_1_-type ATPase (48), there is no evidence of oxidative phosphorylation for an energy generation in aerobically growing *Z. mobilis* cells (8). The aerobic respiration is involved in other functions in *Z. mobilis* (49–51). *Z. mobilis* produces energy solely through substrate phosphorylation by the ED pathway, by stoichiometry of one ATP molecule synthesis per one glucose consumption.

Moreover, glucose is passively transported into *Z. mobilis* cells by the glucose facilitator proteins without an energy requirement (8, 52). This unusual transportation of glucose allows *Z. mobilis* to uptake glucose approximately 8 times faster than actively growing *E. coli* does (53). This high flux of the ED pathway, known as ‘catabolic highway’, coupled with the active pyruvate decarboxylase and the truncated TCA cycle, allocates only less than 5 % of carbon substrate for building biomass in *Z. mobilis* cells (13, 52, 54). In such a bacterium with the rapid catabolism and the low-efficiency in anabolism and energy production, expansive growth and frequent, accurate division do not appear to be a master plan for surviving in nature.

Our present work shows that *Z. mobilis* spatiotemporal cellular organization is distinctively different from those found in the model bacterial species, where active and reproducible propagations are the main drivers. This study provides evidence that *Z. mobilis* cells favour heterogeneity in their division sites, cell sizes, growth rates and DNA contents among individual cells, under laboratory nutrient-rich anaerobic growth conditions.

The observed heterogeneity resembles a bacterial bet-hedging strategy of preparation for unpredictable environmental stress (55, 56). In fact, *Z. mobilis* cells are tolerant in a broad range of pH, high ethanol content (13), and a high dose of different antibiotics (36). It is tempting to speculate that the robust resilience to certain stress in *Z. mobilis*, likely facilitated by its cellular heterogeneity, is conceptually compatible with its main biological activity, the rapid production of ethanol. Both strategies do not prioritize growth, but rather invest in available nutrients in the defence and preparation for fluctuating environments and competitors. Further investigations on the enigmatic ecology of *Z. mobilis* would shed light on the implication of its cellular heterogeneity (57).

How *Z. mobilis* achieves its heterogenetic phenotype at molecular and cellular level deserves further elucidation, especially on FtsZ behaviour. Intriguingly, FtsZ spatial regulation contributes to variant *Z. mobilis* cell size at birth (Fig. 1C), while temporal FtsZ regulation works in reduction of cell size variance (Fig. 2A and Fig. S3). Notably, more than 50 % of total soluble protein contents in *Z. mobilis* cell are enzymes responsible for the fermentation (8, 58). Considering the simplicity and quantity of metabolism in *Z. mobilis*, particular ED-intermediate metabolites or related enzymes might directly bind to cell-structural proteins to dictate cell organization in *Z. mobilis* cells. Direct connections of particular metabolites to morphogenic proteins were previously observed in other bacteria (59–61). Similarly, organization of multiple chromosomes might be directly interfered by the metabolic process since DNA could function as a phosphorous storage polymer in polyploids (12).

The consequence of heterogeneous DNA crowdedness in Zm6 cells was not implicated in its growth, as there was no correlation between DNA density and cellular size that linearly correlates with cell elongation rates (Fig. 1D and Fig. 5D). One of the advantages of possessing bacterial polyploidy is to confer a resilience to DNA damaging events (62). For example, *Deinococcus radiodurans* can withstand high doses of ionizing radiation, utilizing its polyploidy for recovering damaged DNA (63, 64). Similarly, the polyploidy in *S. elongatus* is important for its resistance against UV radiation (41). Alongside with this line, we hypothesize that the polyploidy might be associated with its unique metabolism in *Z. mobilis*. Under oxic conditions, *Z. mobilis* ETS withdraws electron donors NADH from reactions of alcohol dehydrogenase, causing lower ethanol synthesis that requires the reducing equivalents (65, 66). This directly leads to an accumulation of intracellular acetaldehyde in the *Z. mobilis* cells. One of the toxicities of acetaldehyde is to cause double-stranded breaks in DNA, which might be more pronounced in cells with dispersed nucleoids without organelle (Fig. 5A). Although acetaldehyde excretion mechanism is still unclear in *Z. mobilis* (67), the polyploidy might be useful in repairing *Z. mobilis* DNA damage induced by intracellular acetaldehyde under aerobic sugar-rich conditions.

While oblique spindle-orientation is an important mechanism for generating complexity in eukaryotic cells (68–70), an orientation of division septum in rod-shaped bacteria has not been addressed. Our literature survey found one observed bacterial diagonal-like division, which is associated with the branching formation in the multicellular cyanobacterium (71). Based on available literature, there is possibly no suggested favourable thermodynamics or physicochemical property of FtsZ ring or other divisome proteins to direct strict-90 degree constriction to long axis of rod bacterial cells (72, 73).

We observed that *Z. mobilis* divisional plane was often slightly leaned from the short axis of cell (Fig. 6), likely due to the lack of strict spatial regulation of FtsZ (Fig. 4). The angled constriction at non mid-cell position might generate a force to bend the cell during division. After division, the attached ‘pseudo-curvature’ is a temporary state, which fits to the mentioned heterogeneous strategy. Considering its fast catabolism with inefficient growth, some *Z. mobilis* cells may maintain an attached-curved state for a long time in their natural habitats.

Bacterial curved shape was previously shown to be vital for some pathogenic bacteria by being associated with motility and colonization during infection (74). Bacterial curved shape was also shown to be implicated in surface-attachments of *C. crescentus* in flowing aquatic environments (75). Since *Z. mobilis* do not attach surface, and 70% of *Z. mobilis* strains, including Zm6, are non-motile (13), this form of curvature might be rather involved in facilitation of outer membrane vesicle formations at septa (76).

Despite available genetic tools being relatively limited, there are reaming, exciting questions on *Z. mobilis* cell biology. How *Z. mobilis* unique metabolism is related to its cellular organization, and the biological significance of its polyploidy are currently the most intriguing to us.

## Material and methods

### Strains, construction and growing condition

The strains and primers used in this study are listed in the table 1. Used medium for growing *Z. mobilis* was a complex medium containing glucose (20 g/L), NH_4_SO_4_ (1 g/L), KH_2_PO_4_ (1 g/L) and MgSO_4_ (0.5 g/L). The *Z. mobilis* cultures (12 ml) were grown in capped test tubes shaken at 200 rpm, under anaerobic condition at 30 °C throughout this study. The *E. coli* culture was grown in the LB medium in baffled flasks shaken at 200 rpm at 37 °C.

**Table 1.**
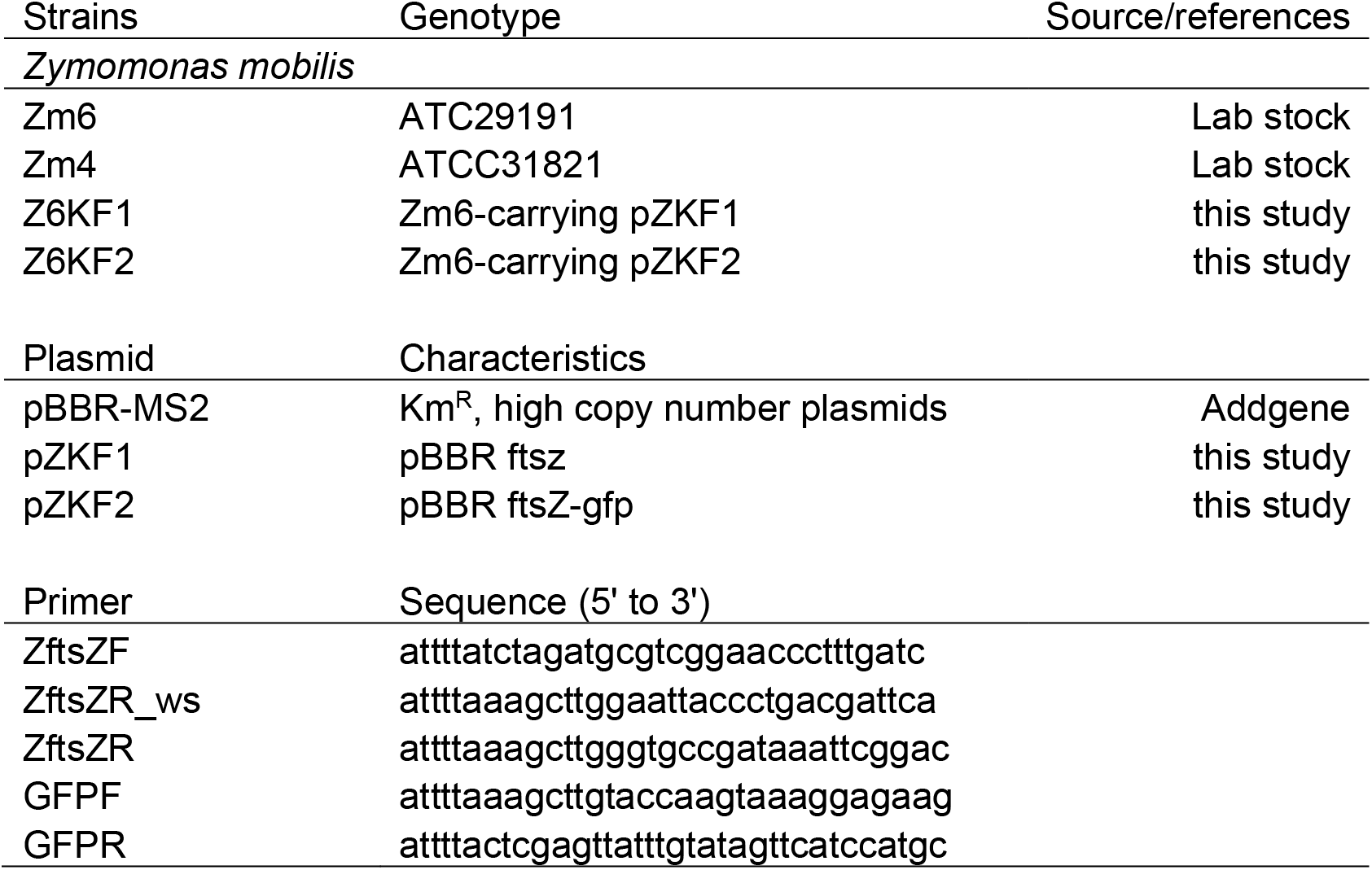
A list of *Z. mobilis* strains, primers and plasmids used in this study.

The plasmid pZKF1, pBBR-ftsZ, was constructed as follows; Zm6 *ftsZ* (ZZ6_0455) was amplified by Phusion polymerase (Thermo Scientific Fisher) using the primers ZftsZF and ZftsZR and genomic DNA of Zm6 as a template. Genomic DNA was extracted combining a lysozyme method and the D-neasy blood tissue kit (Qiagen). The product was purified by the PCR purification kit (Qiagen) and cut by XbaI and HindIII (New England Biolabs). The plasmid pBBR-MS2 was digested with XbaI and HindIII. The cut plasmid and PCR product were then purified and ligated by T4 ligation (New England Biolabs). For constructing the pZKF2 (pBBR-ftsZGFP), Zm6 *ftsZ* was amplified using the primers ZftsZF and ZftsZR_ws. The product was purified and digested with XbaI and HindIII. The *gfpmut3* was amplified from our lab plasmid pSB-M1g with the primers GFPF and GFPR by Phusion polymerase. The product was purified and digested with XbaI and XhoI. The plasmid pBBR-MS2 was cut with XhoI and HindIII. All the products were purified and ligated together by T4 ligase, constructing the pZKF2 carrying *ftsZ*-*gfp* with native *ftsZ* upstream promoter. Sanger sequencing (Eurofins) confirmed that all the constructions was correctly made. Transformation to *Z. mobilis* was done by an electroporation (2.5 kV) using disposable electroporation cuvettes with 0.2 cm gap (VWR) and MicroPulser− (BioRad). The transformants were selected by kanamycin (300 μg/mL).

To observe FtsZ-GFP localizations in Z6KF2 cells, the Z6KF2 culture was grown in the medium with kanamycin (300 μg/mL) overnight. The fully-grown cultures were then sub-cultured in the medium without kanamycin (start OD_600_ 0.05) and grown until its OD_600_ became 0.4 – 0.8. The cells were then mounted on the complex-medium agarose pad (1% w/v) for imaging. According to our pilot experiments, the plasmid stability appeared to be stable without selection marker for at least 30 generations.

For the salt-induced filamentation study, growing Z6KF2 cultures in the regular complex medium with kanamycin (300 μg/mL) were sub-cultured to the complex medium supplemented with additional NaCl 0.2M but without kanamycin. The cultures were grown for 5 hours, and the cells were transferred to the agar-pad of the identical medium without NaCl and imaging was performed.

### Light microscopy

Zeiss Axio Imager Z2 microscope (ZEISS) was used with Axiocam MR R3 (ZEISS) camera for capturing fluorescence or phase contrast imaging. Nikon Ti Eclipse epifluorescence microscope (Nikon) equipped with Hamamatsu C11440-22C camera was used to capture time-lapse imaging. Microscope objectives Nikon 60x Oil (NA1.4) Plan Apo was used for imaging the cells.

For capturing phase contrast images of Zm6 cells, the growing culture of Zm6 was directly mounted on the agarose pad (1% w/v) of an identical medium and sealed by a cover glass for imaging. This was basically applied to time-lapse imaging as well. The cells were incubated at the sealed state at room temperature for about 30 minutes before time-lapse imaging.

Membrane staining; FM4-64 (20 μg/ml) was added to the growing culture, and the cells were incubated for 15 minutes at room temperature. The stained cells were washed by phosphate-buffered saline (PBS) once before imaging. Fluorescent images were analysed by ZEN pro software (ZEISS). Statistical significance was calculated using sigmaplot (Systat Software).

DNA staining; actively growing Zm6 culture was incubated with SYTO™ 9 Green Fluorescent Nucleic Acid Stain (Thermo Fisher Scientific) at a concentration of 0.5 or 11 μM for 10 minutes, or, with DAPI, 4’,6-Diamidino-2-Phenylindole, Dihydrochloride (Thermo Fisher Scientific) at a concentration of 50 μg/mL for 20 minutes at room temperature. The stained cells were washed in PBS before mounting on PBS agarose pad for imaging. We also imaged stained cells mounted on the agarose pad made of complex medium used for liquid culture, and observed similar results using PBS agarose pad. For fixations of cells, *Z. mobilis* cells were fixed by an 10% formalin (v/v) for half hour. The fixed cells were washed by PBS for 3 times before imaging. For visualizing *E. coli* DNA, the growing cells in LB supplemented with glucose 0.1% was stained by DAPI at a concentration of 5 μg/mL for 15 minutes at room temperature. The stained cells were transferred to PBS agarose pad for imaging.

To visualize growing site in *Z. mobilis*, HADA (Tocris) was used at concentration of 500 μM. Zm6 cultures were incubated with HADA at 30 °C for 20 minutes. The stained cells were washed by PBS for twice and mounted on PBS-agarose pad for imaging.

### Image analysis

Cell length was used as a proxy for cell volume in *Z. mobilis* throughout the study. In figures 1A and 1B, cell length and width were measured using Zen pro software. In figure 1D, Elongated cell length was measured using Image J (NIH) and ZenPro. Several cells started to constrict and slightly curved during the 40 minutes of measurement during the time-lapse imaging. In this case, long axis of two cell compartments were separately measured and the combined length was measured for calculating elongated cell length in the figure 1D.

In figure 2, the time-lapse imaging was analysed using Image J. Cell cycle time was defined as time between completions of division, and the completion of division was judged by visible line of septa between two cells (Fig. 2A and 2B). Our time-lapse imaging was time-limited due to the employed agar pad method, and the several small cells (cell length < 1.5 μm) that were generated during imaging did not complete division during the whole time-lapse imaging, and thus these were not included in the analysis (Fig. 2A and 2B).

In figures 3 and 5, Fluorescence (FL) signal intensity profile was measured using profile function of software Zen 2.3 pro. Estimation of background fluorescence signals was done by measuring FL intensity of cell free backgrounds close to measured cells. The measured FL intensity of FtsZ-GFP and nucleoid was subtracted by the background FL intensity for the plot and FL profiles (Fig. 3, 5 and S3). In the figure 5C and 5D, the data set from Zm6 cells stained with 0.5 μM Syto9 was used to obtain the plot.

For measurement of FL intensities within single cell, the measured cells were picked up in the phase contrast images, in order to avoid possible biasing (Fig. 5B-D and Fig. S6). FL intensities at all measured points over long cell axis was averaged to calculate the averaged FL intensity (Fig. 5D). The data set was normalized by the highest intensity being 1. The averaged FL intensity was interpreted as crowdedness of DNA, due to dispersed nucleoid localization in Zm6 cells. For calculating an approximate abundance of DNA per single cell compartment in Zm6 cells, the normalized FL intensity was multiplied by the cell length (Fig. 5E). Visualization of nucleoids in the strain Z6KF2 was performed with staining DAPI at 30 μg/mL for 15 minutes and washing by PBS before mounting on the complex medium-agarose pad for imaging (Fig. 5F).

### Electron microscopy

Three microliters of mid-exponential phase *Z. mobilis* Zm4 or Zm6 cells were applied to glow-discharged holey carbon grids (Quantifoil R3.5/1 Cu 200-mesh) and plunge-frozen in liquid ethane using a Vitrobot Mark IV (Thermo Fisher Scientific). Image acquisition was performed using a Titan Krios transmission electron microscope (Thermo Fisher Scientific) operated at liquid nitrogen temperature at 300 keV and equipped with a Falcon 3 direct electron detector (Thermo Fisher Scientific). Cells were imaged at 11,000x magnification in low dose mode with 1-2s exposure.

## Acknowledgements

The authors gratefully acknowledge Elina Balodite and Uldis Kalnenieks for their advices on handling *Z. mobilis* culture and for donating the plasmid pBBR-MS2. The authors also would like to thank Rhami Lale for a donation of the plasmid pSB-M1g. The study was supported by Norges Forskningsråd (grant number 258657).

